# Temporal resolution of single photon responses in primate rod photoreceptors and limits imposed by cellular noise

**DOI:** 10.1101/421263

**Authors:** Greg D. Field, Valerie Uzzell, E.J. Chichilnisky, Fred Rieke

## Abstract

Sensory receptor noise corrupts sensory signals, contributing to imperfect perception and dictating central processing strategies. For example, noise in rod phototransduction limits our ability to detect light and minimizing the impact of this noise requires precisely tuned nonlinear processing by the retina. But detection sensitivity is only one aspect of night vision: prompt and accurate behavior also requires that rods reliably encode the timing of photon arrivals. We show here that the temporal resolution of responses of primate rods is much finer than the duration of the light response and identify the key limiting sources of transduction noise. We also find that the thermal activation rate of rhodopsin is lower than previous estimates, implying that other noise sources are more important than previously appreciated. A model of rod single-photon responses reveals that the limiting noise relevant for behavior depends critically on how rod signals are pooled by downstream neurons.

## Introduction

Sensory receptors provide us with the raw electrical signals that we use to learn about our environment. Because of this, transduction noise – i.e. variability in the magnitude, latency and kinetics of these signals – limits the reliability with which an organism can determine the properties of external stimuli. Thus, the relationship between sensory performance and the limits imposed by transduction noise reveals the efficiency of sensory processing and provides clues about the underlying mechanisms [2-6].

These considerations have strongly shaped the investigation of rod-mediated vision and the mechanisms that support it. More than a century of work shows that dark-adapted human observers can detect weak flashes of light with a sensitivity approaching the limits set by noise in rod photoreceptors [7]. Past work, however, has focused on the ability to detect photons while neglecting the accuracy with which the time of photon absorption is encoded. Imprecisely signaling the timing of photon absorptions will limit the ability of downstream circuits to perform critical tasks such as determining the direction and speed at which objects in the environment are moving [8, 9]. Furthermore, more accurate measures of rod noise are needed to determine if they indeed limit behavioral detection sensitivity and to understand their impact on limiting temporal precision [3].

Noise in rod phototransduction consists of three main sources: (1) thermal isomerization of rhodopsin [10, 11]; (2) continuous noise produced by spontaneous activation of phosphodiesterase (PDE) [10, 12]; and (3) single photon response variability produced by trial-to-trial fluctuations[13] in the active lifetime of rhodopsin [1, 14-17]. Each of these noise sources has a distinct effect on the photocurrent. For example, thermal isomerizations are rare, but indistinguishable from a light response while continuous noise has a smaller amplitude than the light response but is omnipresent. Thus, these noise sources are likely to place different limits on detection sensitivity versus temporal precision. Furthermore, the relative importance of these noise sources depends on how rod signals are pooled in downstream neural circuitry. For example, thermal activation of rhodopsin is a rare in a small pool of rods but becomes frequent in a pool of 10,000 rods (similar to the receptive field of some ganglion cells). Thus, understanding how transduction limits rod vision requires a consideration of downstream computations.

Here, we show that the temporal resolution of the single photon response is ∼50 ms, much shorter than the ∼750 ms response duration or the ∼250 ms assumed integration time of dark-adapted vision [18]. We incorporate our measurements of rod noise into a model that reveals the impact of each noise source on detection and temporal sensitivity for individual and pooled rod responses. Temporal resolution is limited primarily by continuous noise with some contribution from fluctuations in the single photon response. For pools of rods relevant for behavior, the limiting source of noise depends on the number of rods, detection versus temporal discrimination tasks, and how rod signals are combined or pooled by readout circuits. Finally, our measurements of rod noise indicate that the thermal activation rate of rhodopsin is about half that of previous estimates [7], suggesting that additional noise sources limit behavioral detection sensitivity.

## Results

Our aim was to understand the relationship between noise and sensitivity of rod responses across a broad range of conditions. The results are organized into four sections: (1) measurements of the ability of primate rods to signal the detection and timing of photon arrivals; (2) measurements of each noise source in primate rods; (3) construction of a model of rod responses that uses our measurements of each noise source to reveal their impact on detection and temporal thresholds; and (4) a determinization of the impact of each noise source on detection and temporal thresholds for pools of rods of a size relevant for behavior. We focus on noise generated in the rod outer segment (the photocurrent) for several reasons. Rod outer segment currents are quite independent in nearby rods due to the lack of voltage dependence of the transduction current [19]. This is unlike rod voltages, which are correlated between rods due to electrical coupling [20, 21]; such correlations complicate characterization of rod noise. Furthermore, rod outer segment currents are amenable to quantitative experimental characterization and provide clear limits to the sensitivity of downstream signals and behavior.

### Measuring detection and temporal sensitivity of primate rods

#### A discrimination task to probe sensitivity

Detection and temporal sensitivity were quantified using a two-alternative forced-choice (2AFC) discrimination task [22-24]. Measured rod outer segment currents were used to discriminate flashes delivered at two possible times. A trial consisted of the response to a single flash presented at either an ‘early’ or ‘late’ time (Figure 1A, blue vs. green); the task requires using the response to decide which stimulus was most likely to have been presented. The key determinants of performance -- flash strength and time offset -- were systematically varied. When the offset between possible flash times is large, the task primarily probes detection sensitivity because identifying the correct stimulus depends only on whether the flash is detected (detection limited performance; Figure 1A, left). When the flash strength is large, the task probes temporal sensitivity because the response to the flash will almost always be detectable and identifying the correct stimulus will depend on the time offset (timing limited performance; Figure 1A, right). Individual trials were classified as arising from the early or late flash using an ideal observer analysis based on Fisher’s linear discriminant (Figure 1B, see Methods). Since stimulus discrimination need not be exclusively detection limited or temporally limited, rod performance was also quantified for intermediate stimulus parameters.

**Figure 1.**
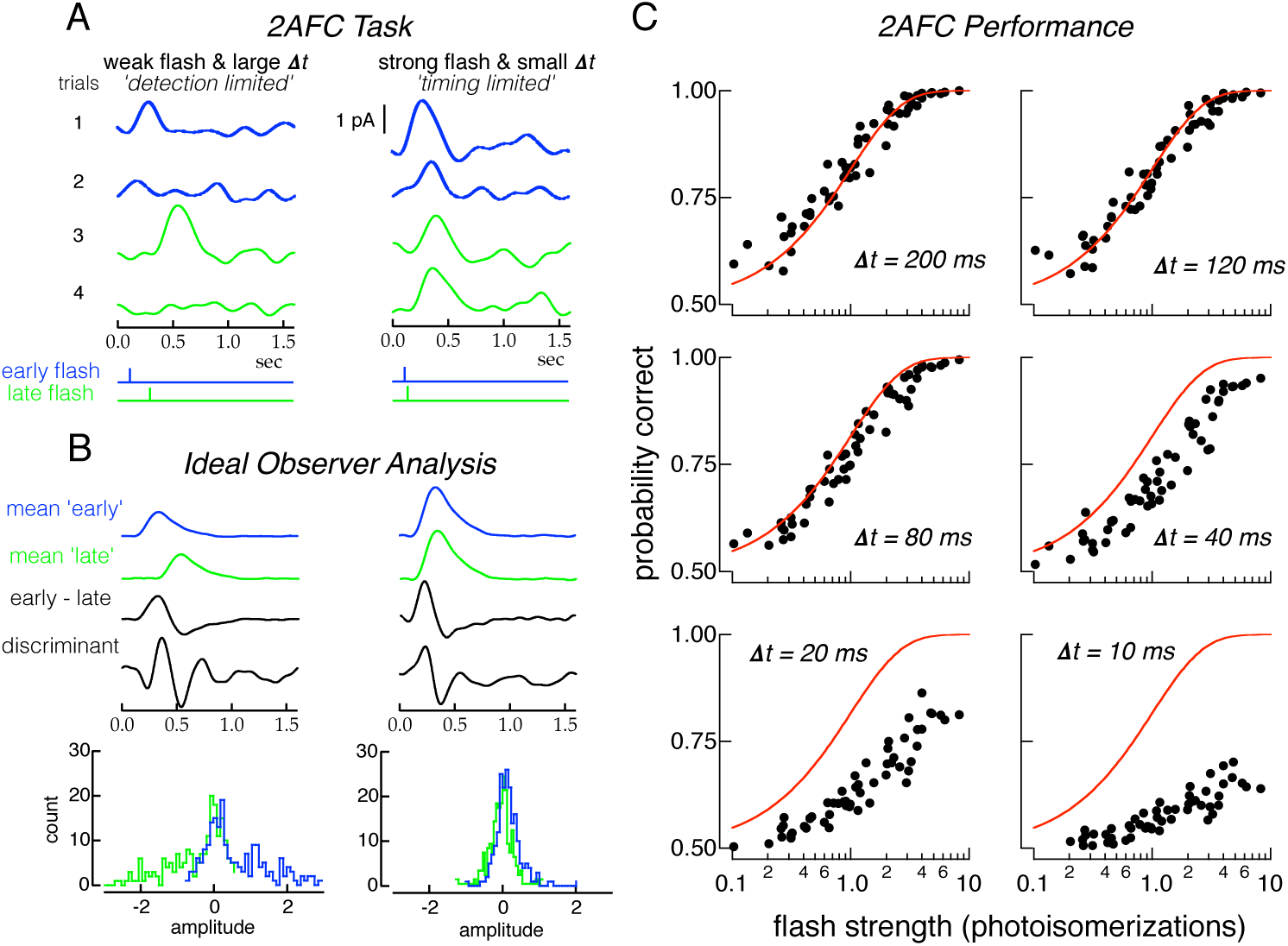
Two-alternative forced-choice (2AFC) task quantifies the detection sensitivity and temporal resolution of rod responses. **A**. Illustration of 2AFC task. *Left:* Detection limited regime of the task. Four responses to a flash producing 0.5 photoisomerizations (Rh*) on average. The flash was delivered at either the early (blue) or late (green) time, separated by 200 ms (bottom stimulus traces). On trials 2 and 4 no response is apparent; on trials 1 and 3 the response clearly identifies the stimulus as ‘early’ or ‘late’. *Right:* Timing limited regime of task. Four responses to a flash producing 3 Rh* on average and a time offset of 10 ms. Responses are evident on every trial but the small time offset and the response variability make classification difficult. **B**. Construction of Fisher’s discriminant used to classify responses. Blue and green show mean responses to ‘early’ and ‘late’ flash times, for detection (left) and timing (right) limited regimes. ‘Early - late’ show difference of the ‘mean early’ and ‘mean late’ responses. Discriminant is Fisher’s linear discriminant (see Methods). Bottom histograms are distributions of inner products between individual trials (A) and the discriminant. Trials to the right (left) of zero would be classified as ‘early’ (late) flashes. **C**. Performance in 2AFC task as a function of flash strength (x-axis in each panel) and time offset *Δt*. Each panel plots probability correct discrimination from 14 rods against flash strength for a *Δt*. Red lines are performance of an ideal photon detector (Equation 1), which is limited only by Poisson variability in photon absorption.

#### Different noise sources limit detection and temporal sensitivity

We used this task and the ideal observer analysis to assess detection and temporal sensitivity of single rods by examining the dependence of discrimination performance on flash strength and time offset. This provides a direct measure of how effectively the time of photon arrivals can be recovered from the slow rod responses. Performance in this task depended strongly on time shift and flash strength (Figure 1C). At large time offsets (120-200 ms), discrimination performance exhibited a transition from chance to perfect over a broad range of flash strengths (Figure 1C, top row). At small time offsets (10-20 ms), discrimination fell short of perfect even for flashes producing 5-10 Rh* on average (Figure 1C, bottom row).

Errors in identifying the correct time of the flash could be introduced by two types of noise: (1) irreducible Poisson variability in the number of photons absorbed from trial-to-trial, and (2) cellular noise in the rod phototransduction cascade. To separate the contributions of cellular and stimulus noise, discrimination performance was compared to that of an ideal photon detector (Figure 1C, red lines). This detector has complete information about the arrival of every photon, and hence its performance is limited only by Poisson variability in the number of photons absorbed from trial to trial. Thus, the ideal photon detector correctly identifies the stimulus, regardless of time offset, on any trial in which one or more photons is absorbed. The performance of the ideal photon detector is given by:

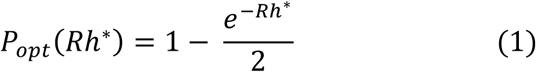

where *Rh*\* is the mean number of photoisomerizations produced by the flash and *e*^−Rh^* is the probability of 0 photoisomerizations given by Poisson statistics. Equation 1 implies that all flashes resulting in 1 or more Rh* and half of those resulting in 0 Rh* are correctly identified. Rod performance cannot exceed that of the ideal photon detector but can fall short due to cellular noise.

Comparing primate rod performance with that of an ideal photon detector (red lines in Figure 1C) indicated that the contributions of stimulus and cellular noise differed for detection and temporal sensitivity. At large time offsets, rod performance matched that of the ideal photon detector (Figure 1C, top). This implies that detection sensitivity is limited by the physical nature of light -- i.e. the division of light into discrete photons and the resulting Poisson variability in the number of photons absorbed from trial to trial (see also [11, 25]). However, at small time offsets, when the task probed temporal sensitivity, rod performance fell short of ideal detector performance. This indicates that cellular noise limits temporal sensitivity.

To summarize rod performance across cells, we plotted probability correct against flash strength and time offset (Figure 2A). Detection sensitivity is represented at one edge of this surface where the time offset is sufficiently large so as not to influence performance (Figure 2A, blue). Detection sensitivity was defined as the flash strength required to achieve 75% correct discrimination, which was 0.7 Rh* (Figure 2B). Temporal sensitivity at a given flash strength can be measured by an orthogonal slice through the discrimination surface (Figure 2A, green line). At a flash strength of 4 Rh* performance was near-perfect at long time offsets, but fell to chance at shorter offsets (Figure 2C). The performance of the ideal photon detector depends only on whether any photons were absorbed, and hence is independent of time offset (Figure 2C, red line). Temporal sensitivity was defined as the time offset required to achieve 75% correct discrimination, which was ∼20 ms for flashes producing 4 Rh*.

**Figure 2.**
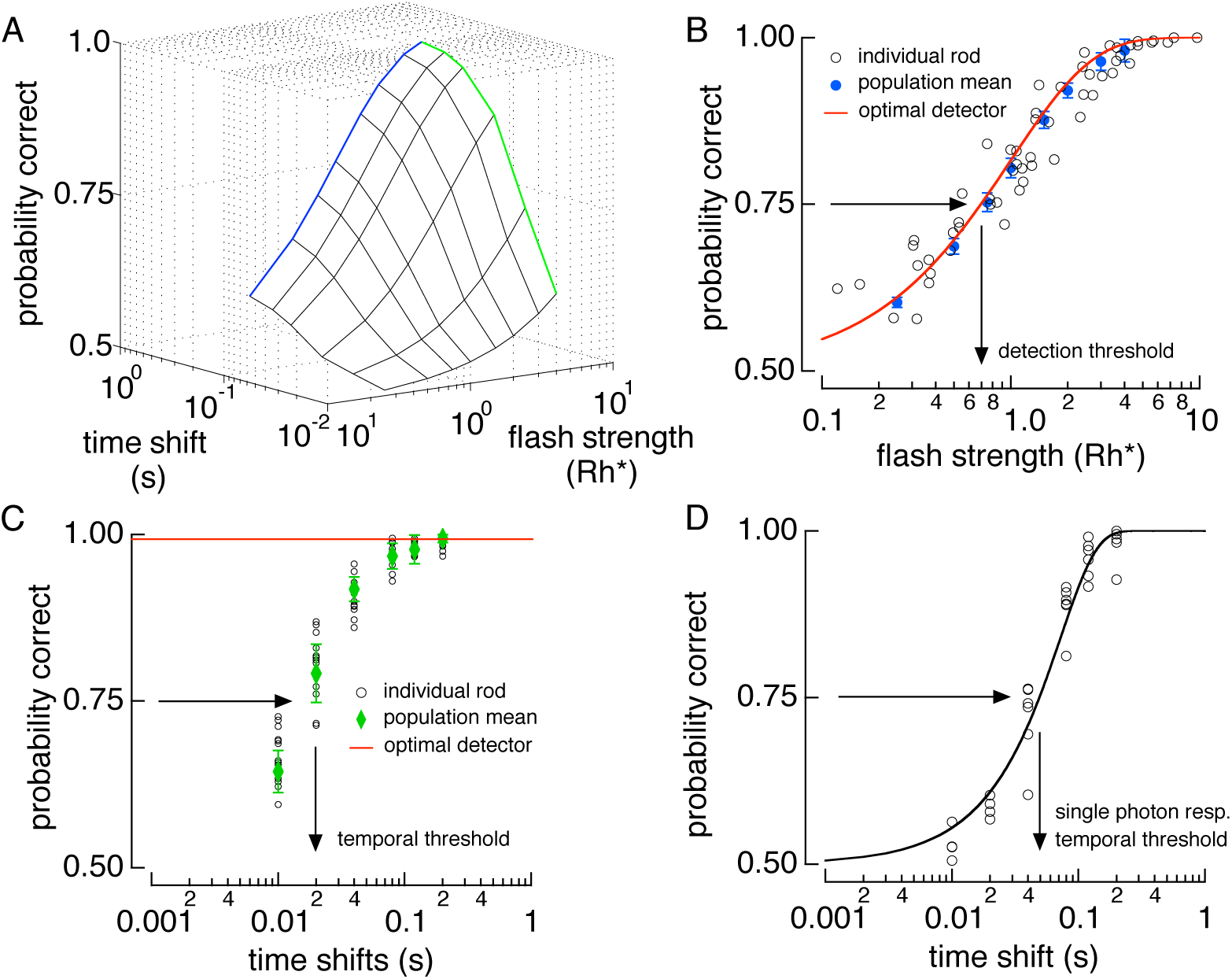
Detection performance is limited by Poisson variability in photon absorptions, while temporal resolution is limited by cellular noise. **A**. 3-D surface of 2AFC performance from rod responses. Blue line is the detection limited side and the green line is the timing limited side. **B**. Detection limited slice of surface in A. Open circles show performance of individual rods at 200 ms time shift. Blue circles show population performance derived from a saturating exponential fit to the data with the error bars representing the 95% confidence interval (see Methods). Red line is the performance of the ideal photon detector (Equation 1). **C**. Timing limited performance in A at a flash strength of 4 Rh*. Population means (green points) are derived from a saturating exponential fit to the rod data (open circles). Red line is the performance of the ideal photon detector. **D**. Temporal sensitivity of the single photon response. Isolated single photon responses were discriminated at different time shifts. Open circles show performance from 6 individual cells. Black line is a cumulative gaussian fit to the data.

Temporal sensitivity defined in this way depends on flash strength: flashes producing more absorbed photons provide more accurate measures of flash timing, as expected from the greater averaging such flashes permit. To provide a more fundamental measure of temporal sensitivity, single photon responses were separated from trials in which 0 or more than one photon were absorbed (see Materials and Methods). This procedure isolated the dependence on cellular noise by removing Poisson variability in photon absorption. Temporal sensitivity of these responses was then measured in a 2AFC task that varied only in time offset (Figure 2D). The time offset required for 75% correct in temporal discrimination was ∼50 ms for single photon responses, considerably less than the ∼750 ms response duration.

What limits the temporal sensitivity of the single photon response? Answering this question required more complete measurements of each source of noise than available from past work. It also required incorporating these measurements into a model to simulate the impact of each noise source on rod sensitivity. We describe our measurements of rod noise first.

### Measuring noise in primate rod phototransduction

#### Noise source 1: thermal activation of rhodopsin

In darkness, the rod photocurrent exhibits occasional large deflections generated by the thermal activation of rhodopsin (Figure 3A) [10, 11, 26, 27]. This noise is indistinguishable from responses generated by photon absorptions (Figure 3A, inset). Previous estimates of the event rate in primate rods have a ∼4-fold uncertainty (95% confidence interval ranges from 0.0027/s to 0.01/s; [11]). To reduce this uncertainty, two procedures were used to estimate the thermal activation rate from long sections of recording in complete darkness (20,980 s total from 13 rods). First, events were counted by eye. This was reliable because the amplitude of these events was high relative to the background noise (e.g. Figure 3A). We counted 78 events, corresponding to a rate of 0.0037 ± 0.0009 events per second (95% confidence limit). Second, the event rate was estimated using a statistical method based on the skew of the distribution of measured current amplitudes [28]. The positive current deflections caused by the thermal activations introduce an asymmetry in the distribution of sampled currents (Figure 3B). Assuming otherwise symmetrical current deflections, the asymmetry can be used to estimate the event rate as

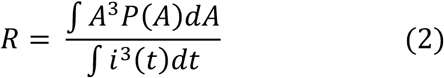

**Figure 3.**
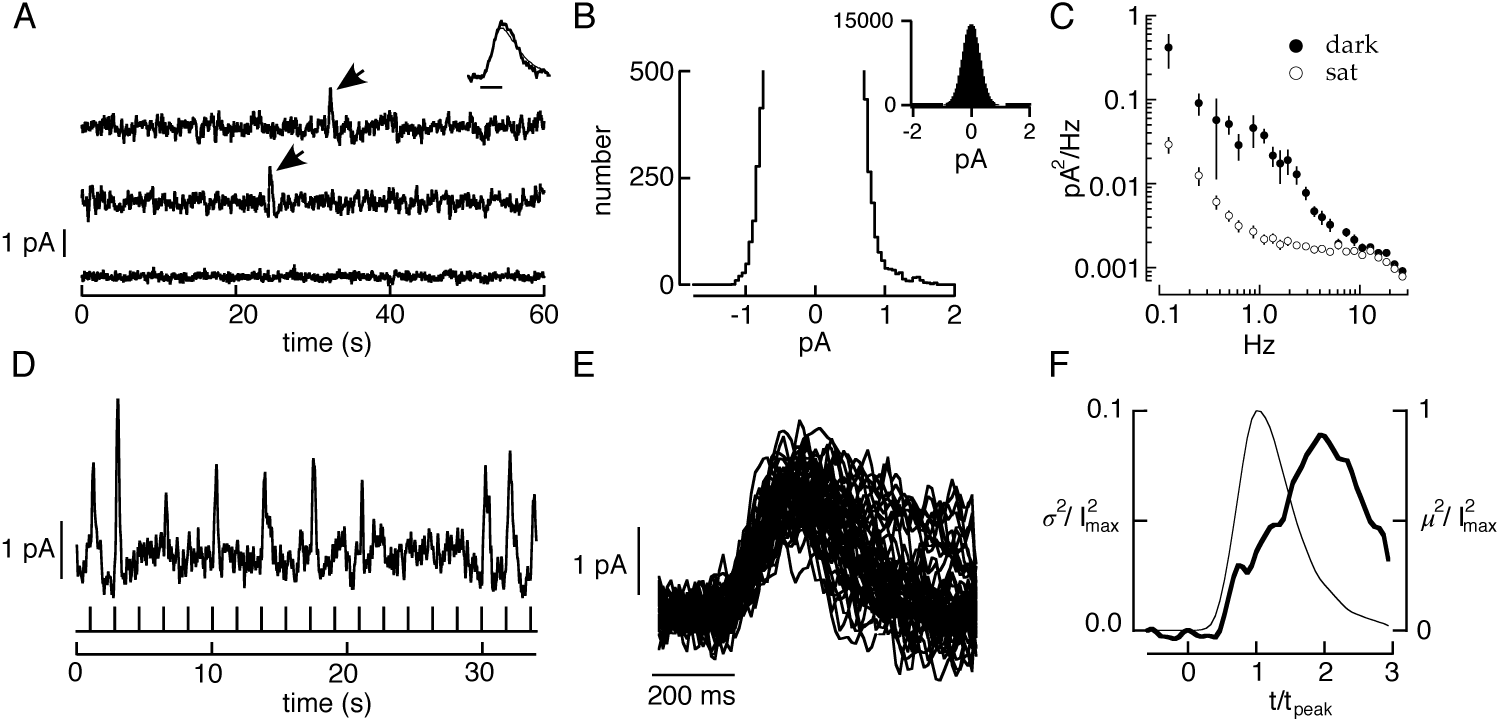
Measurements of three primary noise sources in primate rods. **A**. Current traces from a suction electrode recording of a primate rod. The top two traces are in darkness. Each large positive current deflection (arrows) originated from spontaneous activation of rhodopsin. The bottom trace is recorded in the presence of a bright light which holds the outer segment ion channels closed, exposing the contribution of instrumental noise. *Inset* compares the average of 8 discrete noise events (thick trace) with the cell’s average single photon response (thin trace). Bandwidth: 0-5 Hz; scale bar 200 ms. **B**. Expanded histogram of currents recorded in darkness. The skew in the distribution toward positive current values was used to estimate the rate of thermal activation from Equation 2. The inset shows the full distribution. **C**. Filled circles: power spectrum of the continuous dark noise in recording segments lacking discrete noise events. Open circles: power spectrum in the presence of saturating light. **D**. Rod responses to a repeated dim flash (0.5 Rh* on average). Bandwidth 0-5 Hz. **E**. Isolated single photon responses. **F**. Change in time-dependent variance attributable to photon absorption (variance of singles minus that of failures) collected across rods. The time-to-peak and peak amplitude of the single photon response in each rod were normalized to one prior to combining results across rods. Data (6 rods, 1077 single photon responses total) from [1].

where *P(A)* is the probability of seeing a current deflection of amplitude *A* and *i(t)* is the waveform of the discrete event caused by the thermal activation of rhodopsin (estimated from the single photon response). Applying Equation 2 to our measured rod photocurrents in darkness, we estimated a rate of 0.0034 ± 0.0008 (95% confidence interval) events per second (a total of 72 events for the same 13 cells). The similarity of the two estimates indicates that the true thermal rate is ∼0.0035 Rh*/rod/s per second, at the low end of the range of values from past work (see Discussion).

#### Noise source 2: continuous noise

Continuous noise in phototransduction originates downstream of rhodopsin [10, 12]. Both cellular and instrumental noise contribute to the fluctuations in the measured photocurrent (Figure 3A). We isolated the contribution of instrumental noise by exposing the cell to a bright light that eliminated the outer segment current (Figure 3A, bottom trace and 3C, open circles) [10]. Assuming that cellular and instrumental noise are independent and additive, cellular noise can be isolated by subtracting the power spectrum of the fluctuations in bright light from that in darkness (Figure 3C). Continuous noise was isolated by restricting this procedure to sections of recording lacking thermal activations of rhodopsin. After subtracting instrumental noise, continuous noise was characterized by the residual power spectrum (Figure 3C). The standard deviation of the continuous cellular noise was ∼20-25% of the peak amplitude of the single photon response, similar to previous estimates in toad and primate [10, 11, 25].

#### Noise source 2: single photon response fluctuations

The rod single photon response exhibits trial-to-trial variability that reflects variability in the active lifetime of a rhodopsin molecule [1, 14, 16, 17, 29-31]. To measure and fully describe this variability, dim flashes producing one Rh* (singles) were isolated from trials producing zero or multiple Rh* (Figure 3D-F; Materials and Methods). Data were combined across cells by normalizing the time-to-peak and peak amplitude of each cell’s average single photon response to unity. Combining data across cells (1077 single photon responses from 6 cells) permitted identification of subtle response variations not apparent from the 100-200 single photon responses from a single cell.

The time-dependent variance of the normalized single photon responses reached a peak well after the mean (Figure 3C), indicating that the shape, not just the amplitude, of these responses varied from one to the next [1, 14, 15]. While the time-dependent variance is frequently used to quantify this variability, it is an incomplete measure. Specifically, it does not specify temporal correlations -- i.e. the variance does not determine whether the deviation of the response at one time is correlated with the deviation at another time. Such temporal correlations are described by the covariance. Here we provide the first measurements of single photon response covariance.

The covariance, when measured at a set of discrete time points, forms a matrix and the eigenvectors of this matrix provide a natural, low dimensional, choice for representing the covariance (Figure 4A-C). The eigenvectors identify the characteristic fluctuations across time (Figure 4C), while the associated eigenvalues identify the relative amount of explained variance (Figure 4B). Thus, the eigenvalues and vectors provide a compact representation of temporal fluctuations in the single photon response. We isolated the change in covariance associated with the single photon response (see Methods). The first 5 eigenvectors of the resulting covariance matrix captured >99.9% (92.34, 5.84, 1.41, 0.19, 0.13%) of the total single photon response variance.

**Figure 4.**
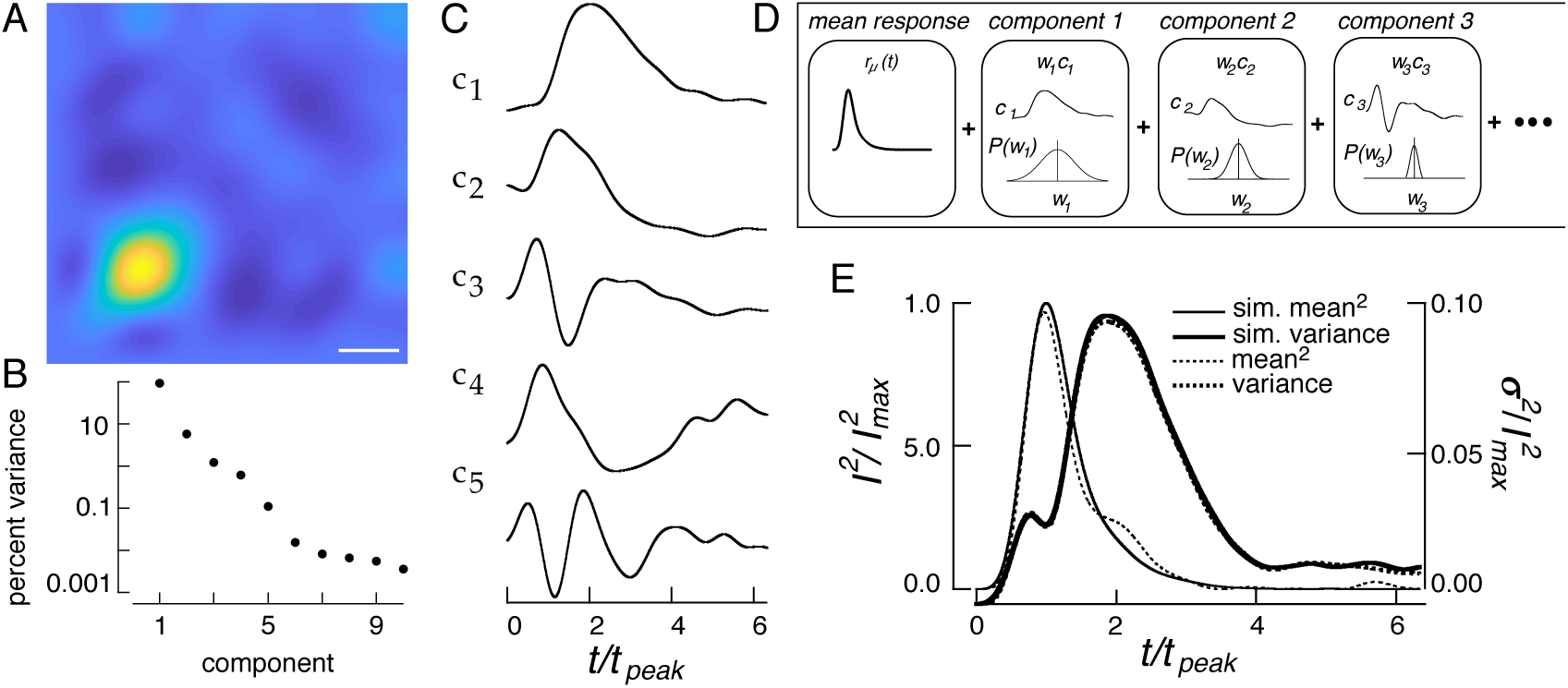
Using the response covariance to simulate single photon responses. **A**. Covariance matrix of the single photon response. Time of flash is the lower left vertex. Scale bar, 300 ms. **B**. Percent of the total variance captured by each eigenvector (c_i_) of the covariance matrix in A. **C**. The five components used to simulate single photon responses. **D**. Generation of response from components, as in Eq. XX. A simulated responses was generated by combining single photon responses with two components of dark noise. Each single photon response was generated as a sum of temporal components corresponding to eigenvectors of the corrected covariance matrix. Each component was weighted by a coefficient drawn from a Gaussian distribution with variance corresponding to the appropriate eigenvalue. **E**. Time-dependent variance and the squared mean single photon response of the model compared with those measured experimentally.

This compact description of single photon response fluctuations is useful because it provides an efficient way for simulating single photon responses and manipulating the magnitude of response fluctuations. Specifically, we express a single photon response *r(t)* as an average response plus a variable contribution from the basis functions, *c*_*i*_*(t)*, given by the eigenvectors of the covariance matrix (Figure 4C):

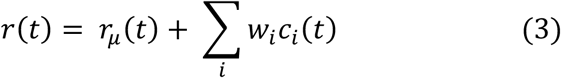

where *r*_*µ*_*(t)* is the mean single photon response, and *w*_*i*_ are weights that describe the contribution of each component *ci* to a particular response (Figure 4D). To illustrate this approach, consider responses that vary only in magnitude. Simulating these responses involves drawing one weight (*w*_*1*_) for each response from an experimentally determined amplitude distribution, multiplying by the average single photon response (*c*_*1*_*(t)*), and adding this product to the mean single photon response (*r*_*µ*_*(t)*). In this special case, the basis function capturing variability equals the mean single photon response (or a scaled version of it) because this is the only ‘direction’ in which the response varies.

To completely capture the fluctuations in single photon response shape, we used the first five eigenvectors of the single-photon response covariance matrix, *c*_*i*_*(t)* (Figure 4C-D). Each response was described by a set of five specific weights *w*_*i*_, applied to the five eigenvectors. The distribution of these weights across responses was well approximated by a zero-mean Gaussian distribution, with a variance given by the eigenvalue associated with that component. This yields a simple procedure for simulating single photon responses: random weights *w*_*i*_ are sampled from appropriate Gaussian distributions, used to scale each of the five eigenvectors, and these scaled eigenvectors are added to the mean single photon response (Figure 4D). This procedure reproduces both the mean squared single photon response and the time dependent variance (Figure 4E).

### A generative model to test the relative importance of different rod noise sources

This section describes the full generative model for rod responses based on their measured noise. The model allows each noise source to be varied independently, thereby allowing a determination of how each noise source impacts detection and temporal sensitivity of individual and pools of rods.

#### Model construction and validation

A simulated rod response was generated by the following equation:

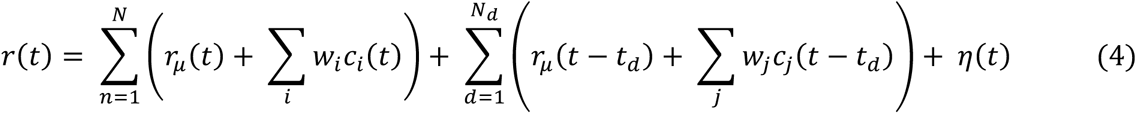

The first term accounts for Poisson variability in the number of photons absorbed by the flash and variability in the resulting single photon responses. The number of absorbed photons, *N*, was drawn from a Poisson distribution with a mean equal to the flash strength, and each single photon response was generated following Equation 3 (Figure 4). The second term accounts for thermal activation of rhodopsin. The number of events, *N*_*d*_, was determined by drawing from a Poisson distribution with a mean equal to the thermal activation rate multiplied by the response duration. Discrete noise events were simulated identically to single photon responses, shifted to occur at a random time *t*_*d*_ relative to the flash, and added to the response. Finally, continuous noise *η(t)* was simulated by filtering Gaussian noise to match the measured continuous noise power spectrum (Figure 3C). The parameters in Equation 4 come directly from experiment with no free parameters.

To validate our model for rod responses, we compared the sensitivity of simulated and recorded responses in the 2AFC task. Simulated responses were generated at ‘early’ or ‘late’ times (as in Figure 1). Model construction and testing used data from different rods to protect against overfitting. The detection and temporal sensitivity of the measured and simulated responses closely matched (Figure 5). The detection sensitivity of simulated rod responses matched that of an ideal photon detector (Figure 5B), and their temporal sensitivity exhibited a threshold at ∼20ms for a flash producing 4 Rh* on average (Figure 5C). Furthermore, the temporal sensitivity of simulated single photon responses matched the data, exhibiting a threshold of ∼50ms (Figure 5D). Thus the simulated responses accurately reproduce the performance of real rods at encoding the arrival and timing of photons.

**Figure 5.**
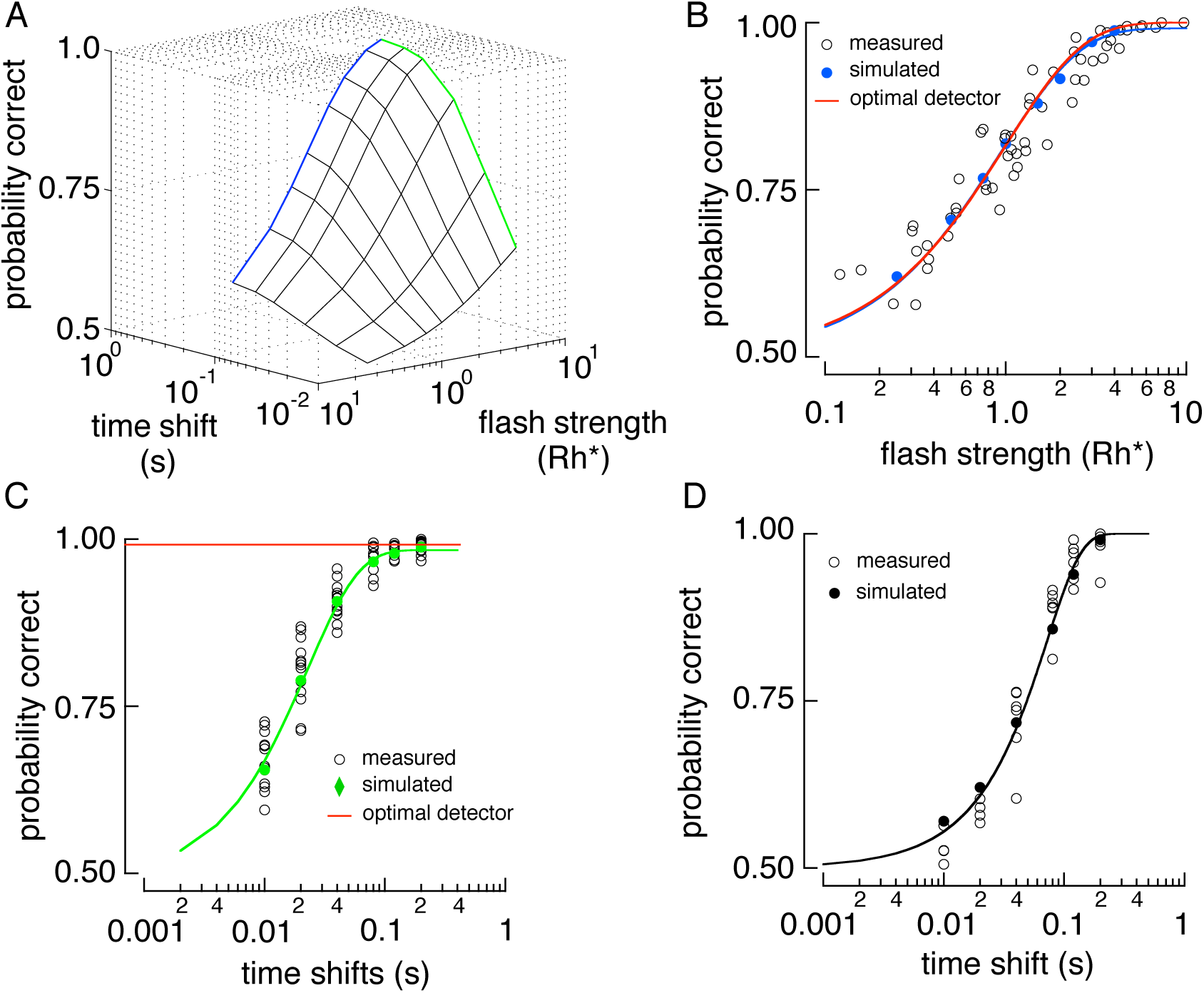
Comparison between the sensitivity of the real and simulated rod responses. **A**. Sensitivity surface of a simulated rod in the 2AFC task of Figure 1. The blue line is the detection limited end of the surface, and the green line is the timing limited end of the surface. **B**. Detection limited slice of the surface in **A**. Closed circles show model performance, and open circles results from Individual rod data replotted from Figure 2. Red line is performance of the ideal photon detector. **C**. Timing limited slice of the surface in **A**. **D**. Temporal sensitivity of simulated and measured single photon responses. Black curve is a cumulative gaussian fit reaching 0.75 probability correct at 50 ms.

#### Controlling response fluctuations is important for detection sensitivity

To test the impact of different rod noise sources on sensitivity, we used the model of rod responses (Equation 4) to vary each noise source independently and used the 2AFC task to measure the detection and temporal sensitivity of the resulting simulated responses (see Methods). Detection sensitivity was robust to changes in rod noise; decreasing any noise source minimally changed performance (Figure 6A, black, green and blue). This is consistent with the observation that detection sensitivity matched that of the ideal photon detector, which is limited only by Poisson variability in photon arrival (Figure 2). Furthermore, increasing any noise source by as much as a factor of 10 produced a relatively small change in detection sensitivity. This is unsurprising for discrete noise events, which occur rarely, and continuous noise, which makes a much smaller contribution to response variance than Poisson fluctuations in photon absorption. Insensitivity to variability in the single photon response may originate because fluctuations are relatively small at the time the response reaches a peak amplitude (Figure 3F) [14]. We tested this hypothesis by simulating single photon responses that varied only in amplitude (i.e. were described by a single component *c*_*1*_*(t)* in Equation 3)) but had the same total variance as the measured responses. In this case, detection threshold rapidly increased as single photon response variability was increased (Figure 6A, red). This indicates that deferring response variance until after the response peak improves detection sensitivity and provides a possible functional advantage to the multi-step shutoff of rhodopsin [14](see Discussion).

**Figure 6.**
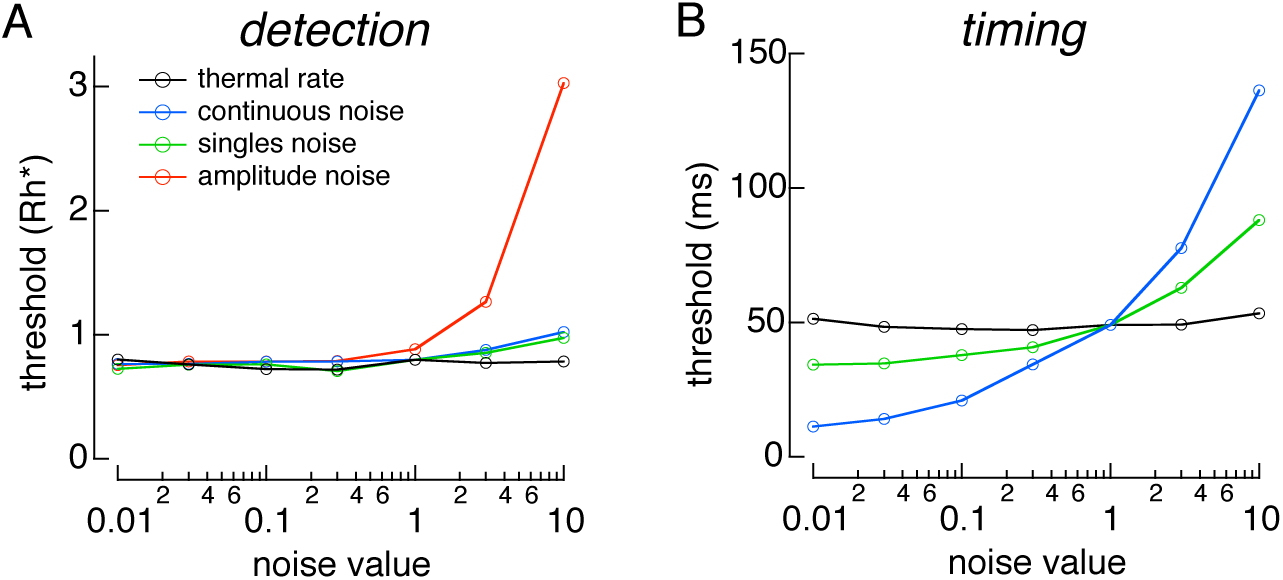
Varying sources of rod noise. **A**. Detection threshold as a function of scaled noise. **B**. Temporal threshold as a function of scaled noise for single photon responses.

#### Reducing response fluctuations and continuous noise improves temporal sensitivity

Temporal sensitivity was probed using a similar approach (Figure 6B). To eliminate the influence of Poisson fluctuations in the number of Rh* produced by a flash, single photon responses were simulated at either an “early” or “late” time, and the response was used to identify the stimulus (as in Figure 2D). Temporal sensitivity was insensitive to the frequency of discrete noise events, again because these events are rare in an individual rod. However, changes in either the continuous noise or the single photon response variability altered temporal sensitivity, with changes in continuous noise having the largest effect. Thus, both noise sources contributed to limiting the temporal resolution of single photon responses in individual rods.

### Impact of rod noise on pooled rod signals depends on linear vs. nonlinear pooling

Behavior is mediated not by single rods, but instead by populations of hundreds or thousands of rods that provide indirect input to retinal ganglion cells [32, 33]. Population size can alter the relative impact of different noise sources as some sources depend on the number of absorbed photons per rod (i.e. Poisson fluctuations in photon absorption and variations in the single photon response), while others do not. We used the model in Equation 3 to quantitatively determine how sensitivity depends on each noise source for the rod populations relevant for behavior, and how sensitivity depends on the ‘read out’ strategy -- i.e. how signals are combined to generate a single output.

We considered pools of 100 and 3,000 rods, which approximate the number of rods that provide (indirect) input to midget and parasol cells at 20º eccentricity. Rod signals were combined either linearly or nonlinearly and used in the our 2AFC discrimination task (see Methods). For nonlinear pooling, the simulated response of each rod was retained if it was more likely a single photon response than noise, and rejected (set to zero) otherwise (see Methods). As described below, the relative importance of different noise sources depended on both the size of the rod pool and the strategy (linear or nonlinear) for integrating signals across rods.

When 100 or 3000 rod signals were linearly pooled, detection thresholds systematically fell short of an ideal photon detector (Figure 7A-B, dashed lines and open circles), unlike the case for single rods (Figure 2). Neither increasing nor decreasing the rate of thermal activation of rhodopsin impacted detection or temporal thresholds at any rod pool size (Figure 7A-D, dashed green lines). Similarly, changing variability in the single photon response modestly impacted temporal thresholds (Figure 7C, dashed black lines), and had no impact on detection threshold (Figure 7A-B). However, increasing or decreasing the amount of continuous noise changed detection and temporal thresholds (Figure 7, dashed blue lines). Decreasing continuous noise 10-fold brought detection thresholds close to the ideal photon detector and nonlinear pooling performance. Thus, continuous noise is the dominant noise source for both detection and timing tasks when rod signals are pooled linearly.

**Figure 7.**
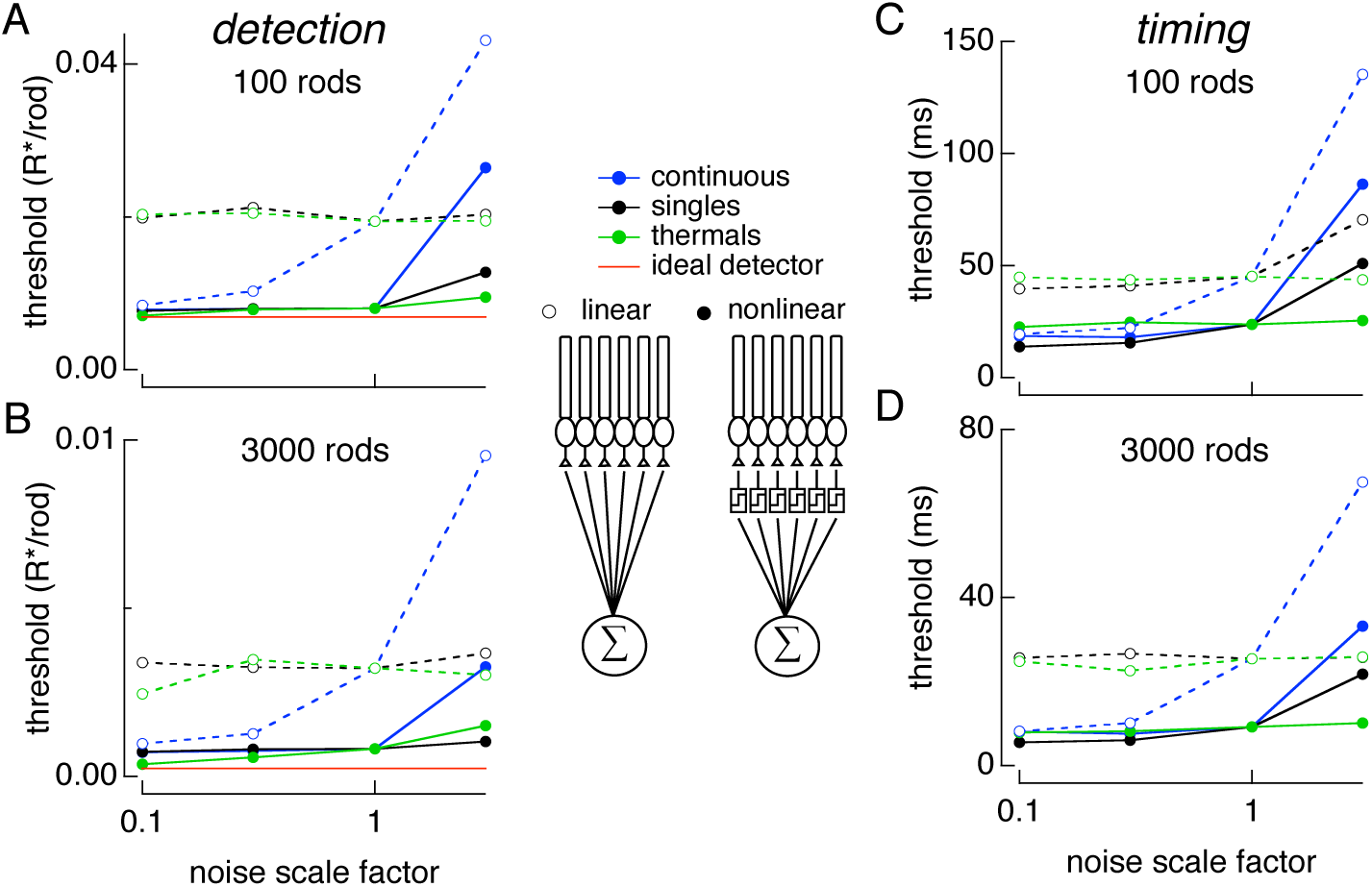
Varying sources of rod noise in pools of rods. Responses of collections of rods were simulated using Equation 4 while varying rod noise sources as in Figure. 6. **A**. Detection threshold for pools of 100 rods as each noise source is varied. Time offset between flashes was 200 ms (detection limited). **B**. Same as **A** for pools for 3000 rods. Results for linear pooling shown with open circles and dashed lines. Results for nonlinear pooling shown with closed circles and solid lines. **C**. Temporal threshold for pools of 100 rods as each noise source is varied. Flash strengths were 0.1 Rh* for linear and nonlinear pooling. These were chosen because each produced >98% correct at large time offsets so that discrimination was timing limited. **D**. Same as **C** for pools of 3000 rods. Flash strengths were 0.01 and 0.003 Rh* for linear and nonlinear pooling respectively. Legend in middle of figure shows schematic for linear and nonlinear pooling.

The relative impact of these noise sources differed considerably when rod signals were pooled nonlinearly. Unlike linear pooling, decreasing the rate of thermal activation of rhodopsin improved detection sensitivity and brought it close to the performance of an ideal photon detector for large rod pools (Figure 7B, solid green line). Thus, with an appropriate nonlinear readout, thermal activation of rhodopsin limited detection sensitivity. However, detection sensitivity remained quite sensitive to increasing continuous noise, reinforcing the importance of regulating the amount of continuous noise (Figure 7A & B, solid blue lines). Interestingly, reducing continuous noise did not improve detection sensitivity, indicating that the amount of continuous noise is matched to the limit imposed by the thermal isomerization rate when an optimal nonlinearity is used to pool rod signals.

Temporal thresholds under nonlinear pooling were minimally changed when increasing or decreasing the thermal activation rate (Figure 7C-D). However, temporal thresholds could be improved by decreasing single photon response variability (solid black lines); reducing continuous noise only improved temporal sensitivity for small pools of rods (≤100). Increasing single photon response variability and continuous noise both increased temporal thresholds, thereby diminishing the temporal resolution of rod vision. The dependence on increases but not decreases in continuous noise reinforces the idea that it is closely matched to the other noise sources when an optimal nonlinear readout is used. As is the case for signals from individual rods, signals from pools of rods permitted recovery of temporal information much finer than the duration of the response itself, or the 200-300 ms integration times often assumed for rod vision (see Discussion).

## Discussion

Past work comparing rod noise with behavioral sensitivity suggests that noise due to the thermal isomerization of rhodopsin limits behavioral thresholds for dim flash detection (reviewed by [3, 34, 35]). If true, the neural circuits that read out rod signals must operate effectively noiselessly - - a considerable constraint. The strength of this inference, however, is limited by at least three issues: (1) experimental uncertainty in the rate of thermal isomerization of rhodopsin in primate rods; (2) lack of consideration of the importance of the other known sources of phototransduction noise; and, (3) a focus on the ability to detect dim lights with little consideration of sensitivity to other aspects of vision (e.g. timing). We consider our work in the context of each of these issues below.

### Thermal isomerization rate lower than previously estimated

Dark-adapted human observers in behavioral detection tasks occasionally report the presence of a flash when no light is delivered. The rate of these false-positive responses suggests an internal noise limiting absolute sensitivity equivalent to ∼0.01 photon-like events/sec/rod [36-38]. These estimates of the noise limiting behavioral sensitivity are close to the estimate of 0.012 events/sec, based on the measured rhodopsin thermal isomerization rate of 0.006 events per sec from primate rods with a 2x correction for the ∼2x larger volume of human rods [11, 35]. However, behavioral and physiological estimates are subject to considerable uncertainty [3, 35, 38]. For example, the 95% confidence interval for the rate of photon-like noise events is 0.0027 to 0.01 events/sec in monkey rods. Thus, a conservative conclusion is that noise from thermal isomerization is within a factor of 2-3 of the noise limiting behavioral sensitivity.

We more precisely estimated the rate of rhodopsin thermal isomerization in monkey rods, placing a tighter bound of 0.0026 to 0.0045 events/sec (95% confidence interval). A recent estimate of the dark light from human behavioral experiments ranged from 0.004 to 0.025 events/sec [34, 38], suggesting that additional sources of neural noise contribute to dark light. An increase in the precision of behavioral measurements will be needed to permit a stringent test of the hypothesis that noise from the thermal isomerization of rhodopsin limits human absolute sensitivity.

### Generative model incorporates all rod noise sources

Three sources contribute to noise in the rod response: thermal isomerization of rhodopsin, continuous current fluctuations, and variability in the single photon response. Past work exploring the implications of rod noise for visual sensitivity has emphasized the importance of thermal isomerizations while largely neglecting the other sources (reviewed by [7]). One reason for this focus is that thermal noise can be easily expressed as a dark light, while other sources of rod noise (or noise downstream of rods) have a more complex relationship to the signal.

To provide a complete picture of how noise limits the fidelity of rod signals, we constructed a parameter-free model incorporating all three noise sources. A key advance in the model was the ability to incorporate accurately the variability in the shape of single photon responses. The model allowed manipulation of different rod noise sources to determine their impact on sensitivity. The model also allowed simulation of signals in collections of hundreds or thousands of rods that form the receptive fields of retinal output cells. While we used this model to investigate the detection and temporal sensitivity of rods, it is a general tool that could be used to probe the encoding of other stimulus features at low light levels.

### Limits to detection and temporal sensitivity: Single rods

The detection sensitivity of single rods approached that of an ideal photon detector limited only by Poisson fluctuations in photon absorption. Temporal sensitivity, however, fell short of that of an ideal photon detector, indicating that it was limited by cellular noise. Nonetheless, temporal sensitivity was much finer than the duration of the response; the single photon response supported a temporal sensitivity of ∼50 ms, about 10-fold less than the duration of the response itself and 2-to-4-fold less than the typically assumed integration time of rod responses [18].

Variability in the single photon response occurs predominantly during response recovery [1, 14, 30]. When single photon response variability was made to (artificially) peak at the time-to-peak of the response, detection sensitivity was degraded (Figure 6A red). Detection sensitivity, however, was insensitive to increasing variability when it occurred late in the response. Thus, the late peak in variability of the single photon response, a signature of inactivation through multiple shutoff steps [1, 14, 15], appears to improve the detection sensitivity of rod signals.

Both continuous noise and single photon response variability contributed to limiting temporal sensitivity. Reducing single photon response variability modestly improved temporal sensitivity, as continuous noise quickly became the limiting factor. Decreasing continuous noise improved temporal sensitivity. However, biophysically, continuous noise arises from instability in PDE, and increasing the stability of PDE would result in less cGMP hydrolysis in darkness and thus a longer duration light response [12]. Hence continuous noise and single photon response variability appear well-matched from the point of view of attaining high temporal sensitivity without prolonging the rod light response (i.e. increasing integration time).

### Limits to detection and temporal sensitivity: Rod populations

Near absolute visual threshold rod signals are conveyed to retinal output neurons --- ganglion cells --- via the specialized rod-bipolar pathway (reviewed by [7, 39]). A key feature of this circuit is that downstream neurons integrate signals from many rods, and this integration can be nonlinear such that single photon responses are preferentially retained, and noise rejected [40]. Other rod pathways in the retina (i.e. rod to Off bipolar cells) appear to integrate rod signals more linearly [40]. We used the generative model to determine how linear and nonlinear pooling of signals across rods influenced the importance of different sources of rod noise.

The detection sensitivity of linearly pooled rod signals depended on the amount of continuous dark noise but was insensitive to changes in the rate of thermal isomerizations. This is because the omnipresent continuous noise quickly dominates the other noise sources as rod signals are added together. The outcome was very different for nonlinear pooling -- the primary purpose of which is to reject the rod signals that only contain continuous noise and retain those that are likely produced by a rhodopsin isomerization, either thermal or photon initiated. Thus, for nonlinear pooling, detection sensitivity was limited by the rate of thermal isomerizations and was insensitive to decreases in continuous noise. Modest increases in continuous noise, however, produced dramatic decreases in sensitivity (increases in threshold in Figure 7). This indicates that continuous noise is just low enough to match the performance limit imposed by the rate of thermal isomerizations when signals are optimally pooled.

Pooled rod responses permitted recovery of the flash timing to a precision < 10 ms - much less than the duration of the rod response and of the ∼200 ms integration time for rod vision [18]. The noise source limiting the temporal sensitivity of pooled rod signals depended on both the number of rods and whether signals were combined linearly or nonlinearity. For linear pooling over small populations of rods (e.g. 100, similar to convergence to a peripheral midget ganglion cell), temporal sensitivity was limited by both single photon response variability and continuous dark noise. However, for larger populations of rods (e.g. 3000, similar to convergence to a peripheral parasol ganglion cell), continuous noise began to limit temporal sensitivity. This switch in the limiting noise source occurs as the fraction of rods absorbing photons at threshold decreases: for larger pools of rods, a smaller fraction need to absorb a photon to reach threshold performance. The smaller fraction of rods absorbing photons means a smaller contribution of single photon response variability relative to continuous noise. With nonlinear pooling, decreasing single photon response variability yielded the greatest improvement in temporal sensitivity. This indicates that limiting single photon response variability is important for maintaining high temporal sensitivity when signals are optimally pooled over a large number of rods.

Pooled rod signals supported temporal sensitivity ∼10 ms even when <1% of the rods absorb photons. Such high temporal sensitivity could support behavioral tasks such as detecting motion at low light levels. Under photopic conditions, temporal resolution approaches 1ms discrimination in apparent motion tasks [41, 42]. Similar behavioral measurements under scotopic conditions would reveal how close the visual system approaches the limits to temporal resolution imposed by rod noise.

These considerations illustrate a more nuanced view of the relationship between rod noise and the sensitivity of downstream cells and of behavior. Specifically, the limiting noise source depends on the task, the number of rods contributing, and how the rod signals are read out. The oft-assumed hypothesis that the rate of thermal isomerization of rhodopsin limits the sensitivity of the rod signals holds under some, but not all, of these conditions.

### Implications for retinal processing and dark-adapted vision

The analysis of the relative importance of different noise sources for different rod pool sizes (Figure 7) shows the benefits of nonlinear pooling. It also sets an upper bound on the sensitivity that can be obtained based on signals in rod outer segments. The pooling models considered, however, neglect several known features of retinal circuitry downstream of the rod outer segment that likely lower the sensitivity of the retinal output signals. For example, gap junctions between rod inner segments could hamper the ability to separate single photon responses from continuous noise [20, 21, 43]. Similarly, downstream cellular and synaptic processes will necessarily introduce noise as signals traverse the retina [3, 39]. Much of such downstream noise is removed by nonlinearities within the retinal circuitry [44]. The importance of downstream noise and processing can be evaluated only in the context of noise in the rod outer segment currents. Hence, we focused here on the constraints imposed by signal and noise at the first step of vision: phototransduction in rod outer segments and the resulting change in photocurrent.

Primate and other mammalian retinas contain ∼30 RGC types [45, 46]. Which and how many of these types participates in vision near absolute threshold remains unclear. However, receptive field size, and thus the number of rod signals pooled by an RGC, varies across eccentricity, RGC type, and across different species. For example, a macaque midget ganglion cell receives convergent input from ∼10 and ∼100 rods at 5° and 20° eccentricity respectively [33]. In contrast, macaque parasol RGCs receive input from ∼100 to ∼3000 rods at these eccentricities. Furthermore, cat alpha cells can receive input from 10,000 to 100,000 rods. Thus, the noise source limiting RGC signal fidelity near absolute threshold is likely to depend on retinal location, cell type, and perhaps species. The brain regions that are responsible for visual perception combine information from multiple RGC types. Thus, future studies aimed at understanding the cortical circuits involved in scotopic vision will be particularly useful in understanding how rod and retinal noise limits behavioral sensitivity.

## Methods

### Photoreceptor recordings

Primate (*Macaca fascicularis* and *Papio anubis*) retinas were provided by the laboratory of D. Dacey at the University of Washington through the Tissue Distribution Program of the Regional Primate Research Center. All procedures followed the guidelines of and were approved by the Administrative Panel on Laboratory Animal Care at the University of Washington. Pieces of retina were obtained in a light adapted state and immediately dark-adapted for >1 hour at 37ºC in bicarbonate-buffered Ames solution equilibrated with 5% CO_2_, 95% O_2_. After dark-adapting, pieces of retina that were not well attached to the pigment epithelium were discarded and the remaining tissue was stored on ice in Hepes-buffered Ames. All procedures after dark adaptation were performed with the use of infrared converters. Rod outer segment currents were recorded with suction electrodes as described previously [1, 47]. The sensitivity, kinetics and continuous noise of rod responses from retina stored in warm bicarbonate-buffered Ames were similar (<15% sensitivity difference, <5% in kinetics, <10% continuous noise SD).

During recording, cells were superfused with bicarbonate-buffered Ames solution warmed to 36.5 - 37.5ºC. Current collected by the suction electrode was amplified, low-pass filtered at 30 Hz (8 pole Bessel) and digitized at 1 kHz. Responses to saturating and half-saturating flashes were measured periodically to check for stability. At the end of a recording, instrumental noise was isolated by exposing the cell to a bright light that eliminated the outer segment current. Only recordings in which cellular dark noise exceeded instrumental noise between 0 and 10 Hz were used in analysis.

Some rods used to measure the thermal activation rate of rhodopsin were loaded with BAPTA to slow the Ca^2+^ kinetics, thereby increasing the signal to noise ratio of the single photon response with respect to the continuous noise. As described previously, a piece of retina was placed in a solution containing 50 µM BAPTA-AM for 20-30 minutes at 37°C before recording [1]. Successful BAPTA incorporation was indicated by biphasic dim flash responses [48].

Light stimulation followed procedures described previously [1]. Briefly, flashes 1-10 ms in duration were delivered using a light-emitting diode (LED) with peak wavelength of 470 nm. Light intensities were measured using a calibrated photodiode (UDT Instruments, San Diego CA). Photon densities (in photons/µm^2^) were converted to photoisomerizations (Rh*) using the collecting area estimated for each rod from the trial-to-trial variability in the responses, assuming that this variability was dominated by Poisson fluctuations in photon absorption.

### Single photon response isolation

Single photon responses were identified and segregated from responses to zero and multiple Rh* by constructing a histogram of response amplitudes. The modes in this histogram corresponded to responses to 0 (centered at zero pA), one (centered at ∼2 pA), and multiple Rh*. Thresholds that identified those responses most likely to be produced by one Rh* were chosen by fitting histograms with a Poisson weighted sum of Gaussians. Control analyses indicated that at least 94% of the isolated responses were indeed single photon responses, with at most 3% contamination from responses to zero absorbed photons and 3% from responses to two absorbed photons [1]. A small number of responses with suspected contamination from thermal noise events were discarded.

The variance or covariance attributable to the single photon response was isolated by subtracting the corresponding measure for responses to 0 Rh*. This assumes that responses to 1 and 0 Rh* make independent and additive contributions to the (co)variance. To check this assumption, we sorted responses to 1 Rh* based on the current immediately preceding the flash. Neither the mean nor variance of the responses depended on the current fluctuation at the time the flash was delivered. Thus any interdependence of the single photon response and continuous noise appeared to be small.

#### Thermal rate measurement

Data were collected in 30 or 60 second records. Reference and saturating flashes were delivered every 2-5 min to insure stable response kinetics and dark current. At the beginning of the recording session and every 10 to 25 minutes thereafter, a series of dim flashes were delivered at one or two flash strengths to estimate the single photon response.

One objective measure of the rate of thermal events relies on the skew of the distribution of measured currents [28] (Equation 2). Three conditions must be met for this approach to yield reliable estimates of the thermal rate: (1) the mean value of the recorded current is zero; (2) the continuous noise is symmetrical; (3) and the single photon response is well estimated. The first condition was met by excluding current records with large amounts of drift (>5 pA) and high-pass filtering the remaining data at 0.1 Hz. This removed low frequency drift in the recording that could cause the mean current to deviate from zero. Occasional large negative electrical artifacts a few ms in duration were excluded by computing the temporal derivative of the current record and searching for threshold crossings >0.2 pA/ms. The current values in a 10 ms window surrounding these events were set to zero. The mean fraction of time set to zero current was 0.22% and did not exceed 0.43% in any cells. Changing the threshold from 0.15 to 0.3 pA/ms decreased the estimated rate of thermal events by <1%. After removing these electrical glitches, data were low-pass filtered with a hard cutoff at 10 Hz. Changing the filter window from 0.1-10 Hz to 0.1-5 Hz increased the estimated thermal event rate by <1%. Stray light rate at the preparation, as measured with a photomultiplier tube, contributed negligibly to the measured rate of thermal rhodopsin activation.

### Discrimination procedures and controls

Much of our analysis relied on using rod responses to discriminate between two possible stimuli. Choosing an optimal or nearly optimal approach to discrimination is critical to identify accurately the detection and temporal thresholds set by rod signal and noise. As described below, Fisher’s linear discriminant provided a near-optimal procedure for the measured rod data.

#### Fisher’s discrimination and dimension reduction

Rod responses were classified as resulting from an early or late flash using Fisher’s linear discriminant [49]. Dimension reduction using PCA was performed on the ensemble of rod responses prior to classification. Ten PCs were used. This number was based on two observations. First, our analysis in Figure 4 indicated that the single photon responses presented at a given time are well captured in a 5-dimensional space. Hence responses presented at two times will be well described in a subspace formed by concatenation of the two spaces, and this 10-dimensional space should provide a complete space for optimal discrimination, though it is possible that fewer than 10 dimensions are required to capture all relevant fluctuations in the responses. Second, discrimination performance reached a plateau at ∼10 dimensions for a number of different discrimination procedures (see below).

After dimension reduction, a single response was removed from the ensemble of responses for a given cell, flash strength and time offset. The remaining responses at two different time shifts were used to calculate Fisher’s discriminant, and the single held out response was classified from its correlation with the discriminant. This procedure was iterated so that each response was summarized by a number, with positive (negative) numbers indicating the response was more likely to be generated by the early (late) flash. The probability correct (ratio of correctly classified trials to all trials) was computed from the test set at each time offset and flash strength.

#### Training set sample size

Discrimination performance can depend on the size of the training set. We checked for such dependence using Fisher’s discriminant with sample sets of 250, 500, and 1000 simulated responses. Performance between 250 and 1000 differed unsystematically by 5.2 ± 9% (mean and SD) averaged across time shifts. A paired student’s t-test and a Wilcoxon rank-sum test did not identify any differences in performance across pairs of thresholds at the same time shift (p = 0.55 and 0.80 respectively). This suggests that discrimination performance was not limited by finite data. Based on these results, training sets for calculating Fisher’s linear discriminant used 500 responses to both early and late flashes, unless otherwise noted.

#### Checking Fisher’s discriminant performance against other parametric discrimination procedures for individual rod responses

Fisher’s linear discriminant is a parametric method for discriminating two multivariate distributions. It is an optimal procedure when the data for both classes are normally distributed with each class having the same covariance [49]. Poisson variability in photon absorption and variability in the rod’s single photon response will cause these assumptions to fail, raising the possibility that other classifiers may exhibit better performance. To check this possibility we tested three additional parametric methods: (1) a difference of means classifier (DM), (2) quadratic discriminant analysis (QDA), (3) model-based cluster analysis (MBCA) [49]. These three classification procedures perform optimal classification with increasingly more complex distributions, but in turn require more parameters to be estimated. In theory MBCA should provide the best performance because our data for each class arise from several multivariate-normal distributions with unconstrained covariance matrices. However, the number of parameters which need to be estimated for such a procedure to perform well is relatively large. Fisher’s discriminant, QDA, and MBCA all performed significantly better (∼30%) than DM at time shifts between 20 and 80 ms and dimensionality >4. However, Fisher’s discriminant, QDA, and MBCA exhibited indistinguishable performance except at the smallest time shifts (5-20 ms) where MBCA consistently performed 10-20% less well.

We also compared the performance of Fisher’s discriminant with a non-parametric classification procedure. In this procedure, responses were divided into training and test data sets. PCA was used to reduce the dimensionality of the data to 10. Variance across each dimension was normalized to one across classes (a whitening transform). An empirical distribution for each flash was created by associating a spherical Gaussian probability density with each point in the training data set. Test data were then classified by choosing the empirical distribution with the higher likelihood for generating the test data. Performance was stable for SDs of the spherical Gaussians between 0.1 and 10 times the median inter-point distance. The performance of this classifier could not be distinguished from that of a Fisher classifier at training sample sets of >250 with a paired student’s t-test or Wilcoxon rank-sum test (p = 0.65 and 0.80 respectively). At training sets of 1000 responses, statistical tests indicated even greater similarity (p = 0.95 and 0.96: t-test, Wilcoxon).

For both parametric and non-parametric discrimination, performance was stable for frequency ranges between 5-10 Hz. Below 5 Hz, the frequency cutoffs impinged on the signal; above 10 Hz, additional instrumental noise contaminated the signal. A frequency range of 0.1-6 Hz was used for discrimination.

### Rod pooling models

To investigate discrimination for pools of rod responses, individual responses were generated using Eq. 4 (see Results). For linear pooling, each rod response in the pool was projected along the discriminant and then summed across all rods in the pool:

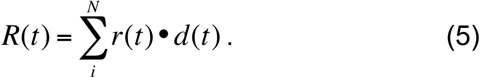

*R(t)* is the summed response, *N* is the number of rods in the pool, *r(t)* is an individual rod response given by Eq. 5, and *d(t)* is the discriminant. If the sum of these projections was greater than zero, the response was classified as being generated by the “early” flash, otherwise it was generated by the “late” flash. If *R(t)* was 0, then the response was randomly classified as an early or late flash with equal probability.

The discriminant, *d(t)*, was learned on a training set consisting of 1000 simulated single photon responses: 500 elicited by the early flash, and 500 by the late flash. A different discriminant was computed for each time offset. Fisher’s discriminant applied to pooled rod responses did not yield performance better than a DM discriminant for any pool size >30 rods or for any set of scaled noise values under linear pooling. This differs from discrimination using responses of single rods (see above), in which case Fisher’s discriminant yielded better performance. This difference likely originates because pooling alters the relative importance of continuous noise and noise due to variability in the single photon response; these two noise sources have quite different covariance structure. Hence for pooled rod responses we used a DM discriminant. Once the discriminant was calculated for a particular pool size and set of noise values, it was used to discriminate 100 novel test trials at each flash time. Threshold performance was defined as the flash strength or temporal offset that yielded 75% correct in the 2AFC task.

For nonlinear pooling, responses were passed through a thresholding nonlinearity [40, 50, 51]. The purpose of this nonlinearity was to suppress continuous dark noise and retain as many single photon responses as possible [52]. The threshold was instantiated by first classifying every individual rod response as more likely to be continuous noise or a single photon response, prior to summing the responses together. The threshold was learned by simulating 500 responses to 0 and 1 Rh* and filtering these in time by the mean single photon response:

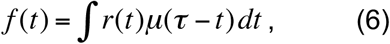

*f(t)* is the filtered response, *r(t)* is the simulated response, and *µ(t)* is the mean single photon response. The maxima of the filtered responses were used to generate an amplitude distribution of responses to 0 and 1 Rh*. The resulting distribution was fit by a sum of two Gaussians (one Gaussian for the 0 Rh* distribution and one Gaussian for 1 Rh* distribution) that were weighted by the Poisson probabilities of observing 0 or 1 photon [40]:

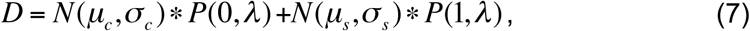

*D* is the distribution of expected response amplitudes, *N* denotes a normalized Gaussian, and *P* is a Poisson probability of observing 0 or 1 photons given a flash strength of *λ*. *µ*_*c*_ and *µ*_*s*_ are the mean of the continuous noise (failures) and singles distributions, *σ*_*c*_ and *σ*_*s*_ are the standard deviations.

The crossing point between the singles and failures distributions was taken as the optimal threshold *T*, for deciding whether a response is more likely a single or a failure. It was computed as the local minimum in D between *µ*_*c*_ and *µ*_*s*_.

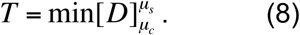

Responses below threshold *T* were more likely to be noise and were multiplied by 0; those above threshold were more likely signal and were multiplied by 1. This procedure set the nonlinear threshold to be matched to the prior probability of photon absorption and the corresponding distributions of signal and noise [40]; nonlinear thresholds were determined separately for each noise condition. This nonlinear thresholding allowed only responses that were most likely true signals to participate in ‘deciding’ whether the flash was early or late.

Thresholded responses were summed and used to generate a discriminant between early and late flash times. The discriminant was generated from 500 trials at both flash times and 100 test trials for each flash time were classified. As with linear pooling, Fisher’s discriminant did not produce significance performance improvements over the simpler DM discrimination, so DM was used.

Theoretically, the optimal instantiation of nonlinear pooling of rod signals is a Bayesian discrimination performed on the pool of rod responses. This differs from the nonlinear pooling approach described above as follows. Rather than using a sharp threshold that multiplies each response by 0 or 1 according to whether the response is more likely a failure or single, every response is weighted by the likelihood ratio of coming from the singles versus failures distributions. Thus, the weights are continuously valued, with small weights associated with responses that are likely noise and larger weights applied to those that are likely single photon responses. The performance of Bayesian pooling and discrimination did not differ significantly from the threshold pooling used and did not change the relative importance of different noise sources as a function of rod pool size or task (data not shown). This is because the likelihood ratio (ratio of probability of single vs failure) changes quite quickly from 0 to 1. We report results based on the threshold nonlinearity for its potentially greater biological relevance [40].

### Manipulating noise in simulated responses

Fluctuations in the single photon response were varied by increasing or decreasing the width of the Gaussian distributions from which the weights, *w*_*i*_, were drawn, continuous dark noise was varied by uniformly scaling its power across frequencies, and the rate of discrete noise events was increased or decreased.

## Acknowledgments

We thank A.P. Sampath, P.H. Li for comments on the manuscript, Dennis Dacey, Matt McMahon and Orin Packer for help with primate tissue, Eric Martinson and Maria McKinley for technical assistance, and Felice Dunn and Thuy Doan for helpful discussions.

Contributions
GDF, VU, EJC, and FR conceived and designed the experiments and analyses. GDF and FR performed the experiments. GDF and FR analyzed the data. GDF and VU generated the rod signal pooling models. GDF and FR wrote the manuscript.

## References

Field, G., and Rieke, F. (2002). Mechanisms regulating variability of the single photon responses of mammalian rod photoreceptors. Neuron 35, 733–747.

Bialek, W. (1987). Physical limits to sensation and perception. Annu Rev Biophys Biophys Chem 16, 455–478.

Field, G., Sampath, A., and Rieke, F. (2005). Retinal processing near absolute threshold: from behavior to mechanism. Ann. Revs Physiol 67.

Faisal, A., and Wolpert, D. (2009). Near optimal combination of sensory and motor uncertainty in time during a naturalistic perception-action task. J Neurophysiol 101, 1901–1912.

Chen, F., Zha, D., Fridberger, A., Zheng, J., Choudhury, N., Jacques, S., Wang, R., Shi, X., and Nuttall, A. (2011). A differentially amplified motion in the ear for near-threshold sound detection. Nat Neurosci 14, 770–774.

Bathellier, B., Steinmann, T., Barth, F., and Casas, J. (2012). Air motion sensing hairs of arthropods detect high frequencies at near-maximal mechanical efficiency. J R Soc Interface 9, 1131–1143.

Field, G., and Sampath, A. (2017). Behavioural and physiological limits to vision in mammals. Philos Trans R Soc Lond B Biol Sci 372.

Gegenfurtner, K., Mayser, H., and Sharpe, L. (2000). Motion perception at scotopic light levels. J Opt Soc Am A Opt Image Sci Vis 17, 1505–1515.

Billino, J., Bremmer, F., and Gegenfurtner, K. (2008). Motion processing at low light levels: Differential effects on the perception of specific motion types. J Vis 8, 14.11-10.

Baylor, D., Matthews, G., and Yau, K. (1980). Two components of electrical dark noise in toad retinal rod outer segments. J Physiol 309, 591–621.

Baylor, D., Nunn, B., and Schnapf, J. (1984). The photocurrent, noise and spectral sensitivity of rods of the monkey Macaca fascicularis. J Physiol 357, 575–607.

Rieke, F., and Baylor, D. (1996). Molecular origin of continuous dark noise in rod photoreceptors. Biophys J 71, 2553–2572.

Lamb, T.D., and Kraft, T.W. (2016). Quantitative modeling of the molecular steps underlying shut-off of rhodopsin activity in rod phototransduction. Mol Vis 22, 674–696.

Rieke, F., and Baylor, D. (1998). Origin of reproducibility in the responses of retinal rods to single photons. Biophys J 75, 1836–1857.

Hamer, R., Nicholas, S., Tranchina, D., Liebman, P., and Lamb, T. (2003). Multiple steps of phosphorylation of activated rhodopsin can account for the reproducibility of vertebrate rod single-photon responses. J Gen Physiol 122, 419–444.

Doan, T., Mendez, A., Detwiler, P., Chen, J., and Rieke, F. (2006). Multiple phosphorylation sites confer reproducibility of the rod’s single-photon responses. Science 313, 530–533.

Gross, O., Pugh, E.J., and Burns, M. (2012). Calcium feedback to cGMP synthesis strongly attenuates single-photon responses driven by long rhodopsin lifetimes. Neuron 76, 370–382.

Conner, J. (1982). The temporal properties of rod vision. J Physiol 332, 139–155.

Baylor, D.A., and Nunn, B.J. (1986). Electrical properties of the light-sensitive conductance of rods of the salamander Ambystoma tigrinum. J Physiol 371, 115–145.

Li, P.H., Verweij, J., Long, J.H., and Schnapf, J.L. (2012). Gap-junctional coupling of mammalian rod photoreceptors and its effect on visual detection. J Neurosci 32, 3552–3562.

Jin, N.G., and Ribelayga, C.P. (2016). Direct Evidence for Daily Plasticity of Electrical Coupling between Rod Photoreceptors in the Mammalian Retina. J Neurosci 36, 178–184.

Geisler, W., Albrecht, D., Salvi, R., and Saunders, S. (1991). Discrimination performance of single neurons: rate and temporal-pattern information. J Neurophysiol 66, 334–362.

Chichilnisky, E., and Rieke, F. (2005). Detection sensitivity and temporal resolution of visual signals near absolute threshold in the salamander retina. J Neurosci 25, 318–330.

Smith, R., and Dhingra, N. (2009). Ideal observer analysis of signal quality in retinal circuits. Prog Retin Eye Res 28, 263–288.

Schneeweis, D., and Schnapf, J. (1995). Photovoltage of rods and cones in the macaque retina. Science 268, 1053–1056.

Luo, D., Yue, W., Ala-Laurila, P., and Yau, K. (2011). Activation of visual pigments by light and heat. Science 332, 1307–1312.

Gozem, S., Schapiro, I., Ferre, N., and Olivucci, M. (2012). The molecular mechanism of thermal noise in rod photoreceptors. Science 337, 1225–1228.

Donner, K., Firsov, M.L., and Govardovskii, V.I. (1990). The frequency of isomerization-like ’dark’ events in rhodopsin and porphyropsin rods of the bull-frog retina. J Physiol 428, 673–692.

Baylor, D., Lamb, T., and Yau, K. (1979). Responses of retinal rods to single photons. J Physiol 288, 613–634.

Whitlock, G., and Lamb, T. (1999). Variability in the time course of single photon responses from toad rods: termination of rhodopsin’s activity. Neuron 23, 337–351.

Caruso, G., Bisegna, P., Andreucci, D., Lenoci, L., Gurevich, V., Hamm, H., and DiBenedetto, E. (2011). Identification of key factors that reduce the variability of the single photon response. Proc Natl Acad Sci U S A 108, 7804–7807.

Sterling, P., Freed, M., and Smith, R. (1988). Architecture of rod and cone circuits to the on-beta ganglion cell. J Neurosci 8, 623–642.

Goodchild, A., Ghosh, K., and Martin, P. (1996). Comparison of photoreceptor spatial density and ganglion cell morphology in the retina of human, macaque monkey, cat, and the marmoset Callithrix jacchus. J Comp Neurol 366, 55–75.

Donner, K. (1992). Noise and the absolute thresholds of cone and rod vision. Vision Res 32, 853–866.

Nelson, P.C. (2016). Old and new results about single-photon sensitivity in human vision. Phys Biol 13, 025001.

Barlow, H. (1956). Retinal noise and absolute threshold. Journal of the Optical Society of America 46, 634–639.

Sakitt, B. (1972). Counting every quantum. J Physiol 223, 131–150.

Koenig, D., and Hofer, H. (2011). The absolute threshold of cone vision. J Vis 11.

Bloomfield, S., and Dacheux, R. (2001). Rod vision: pathways and processing in the mammalian retina. Prog Retin Eye Res 20, 351–384.

Field, G., and Rieke, F. (2002). Nonlinear signal transfer from mouse rods to bipolar cells and implications for visual sensitivity. Neuron 34, 773–785.

Westheimer, G., and McKee, S.P. (1977). Perception of temporal order in adjacent visual stimuli. Vision Res 17, 887–892.

Westheimer, G. (1999). Discrimination of short time intervals by the human observer. Exp Brain Res 129, 121–126.

Hornstein, E., Verweij, J., Li, P., and Schnapf, J. (2005). Gap-junctional coupling and absolute sensitivity of photoreceptors in macaque retina. J Neurosci 25, 11201–11209.

Ala-Laurila, P., and Rieke, F. (2014). Coincidence detection of single-photon responses in the inner retina at the sensitivity limit of vision. Curr Biol 24, 2888–2898.

Field, G., and Chichilnisky, E. (2007). Information processing in the primate retina: circuitry and coding. Annu Rev Neurosci 30, 1–30.

Sanes, J., and Masland, R. (2015). The types of retinal ganglion cells: current status and implications for neuronal classification. Annu Rev Neurosci 38, 221–246.

Baylor, D., Lamb, T., and Yau, K. (1979). The membrane current of single rod outer segments. J Physiol 288, 589–611.

Matthews, H. (1991). Incorporation of chelator into guinea-pig rods shows that calcium mediates mammalian photoreceptor light adaptation. J Physiol 436, 93–105.

Duda, R., Hart, P., and Stork, D. (2001). Pattern classification, (New York: Wiley & Sons).

Berntson, A., Smith, R., and Taylor, W. (2004). Transmission of single photon signals through a binary synapse in the mammalian retina. Vis Neurosci 21, 693–702.

Okawa, H., Miyagishima, K., Arman, A., Hurley, J., Field, G., and Sampath, A. (2010). Optimal processing of photoreceptor signals is required to maximize behavioural sensitivity. J Physiol 588, 1947–1960.

van Rossum, M., and Smith, R. (1998). Noise removal at the rod synapse of mammalian retina. Vis Neurosci 15, 809–821.

